# Thiamine transporter 2 and Janus kinase 2 inhibitor, fedratinib suppresses thermogenic activation of human neck area-derived adipocytes

**DOI:** 10.1101/2025.11.24.690155

**Authors:** Gyath Karadsheh, Emília Kovács, Rahaf Alrifai, Mizuki Seo, Ferenc Győry, Renáta Csatári-Kovács, Éva Csősz, Szilárd Póliska, László Fésüs, Rini Arianti, Endre Kristóf

**Affiliations:** Laboratory of Cell Biochemistry, Department of Biochemistry and Molecular Biology, Faculty of Medicine, University of Debrecen, H-4032 Debrecen, Hungary; Doctoral School of Molecular Cellular and Immune Biology, University of Debrecen, H-4032 Debrecen, Hungary; Department of Surgery, Faculty of Medicine, University of Debrecen, H-4032 Debrecen, Hungary; Proteomics Core Facility, Department of Biochemistry and Molecular Biology, Faculty of Medicine, University of Debrecen, H-4032 Debrecen, Hungary; Genomic Medicine and Bioinformatics Core Facility, Department of Biochemistry and Molecular Biology, Faculty of Medicine, University of Debrecen, H-4032 Debrecen, Hungary

**Keywords:** human adipocytes, thiamine, SLC19A3, fedratinib, JAK, thermogenesis

## Abstract

**Introduction:** Brown adipocytes consume higher amounts of metabolic substrates and regulators including thiamine during adrenergic stimulation supporting heat generation. Our previous findings showed that fedratinib, a potent inhibitor of thiamine transporter (ThTr) 2 and Janus kinase 2 (JAK2), reduced thermogenic activity; however, the underlying molecular mechanisms remain elusive.

**Methods:** Primary human subcutaneous (SC) and deep neck (DN)-derived adipocytes were treated with dibutyryl (db)-cAMP, fedratinib, or the combination of the two compounds after differentiation. Global transcriptomic analysis was performed by bulk RNA-sequencing. Differentially expressed genes were subjected to pathway enrichment analysis. We also utilized publicly available single-cell RNA-sequencing datasets and adiposetissue.org to correlate ThTr2 expression in adipose tissue to clinical parameters of patient cohorts. Amino acid flux was measured by metabolomics.

**Results and discussion:** ThTr2 expression was observed exclusively enriched in the adipocytes cluster within human brown and white adipose tissue. In response to ThTr2 inhibition, the db-cAMP-stimulated upregulation of the canonical thermogenic markers and proton leak respiration, which associates with UCP1-dependent heat generation, was prevented in both adipocyte types. RNA-sequencing found 40 and 41 downregulated genes potentially underlying the metabolic changes in SC and DN-derived adipocytes, respectively, which were involved in various biological pathways, including transcriptional regulation of brown and beige adipocytes differentiation, signaling by interleukins, nicotinamide salvaging, and gene and protein expression by JAK/STAT signaling after interleukin-12 stimulation. The expression of recently identified thermogenesis regulators, such as transglutaminase (TGM) 2 and inhibitor of DNA binding (ID) 1, was also abrogated by ThTr2 inhibition during adrenergic stimulation. Intriguingly, glutamate transporter (GLT) 1 and L-amino acid transporter (LAT) 2 expression was also attenuated by fedratinib, restricting amino acid consumption. Finally, we found that the expression of ThTr2 in human white adipose tissue was inversely correlated with body mass index, waist-hip ratio, leptin secretion, and plasma insulin, glucose, cholesterol and triacylglycerol levels, supporting the importance of thiamine metabolism in adipocyte and metabolic health.

- Fedratinib prevented adrenergic-stimulated upregulation of thermogenic genes.
- The consumption of amino acids and the expression of their transporters decreased by fedratinib during adrenergic stimulation.
- Oxygen consumption and proton leak respiration reflecting heat generation were abrogated by fedratinib.

## 1 Introduction

Adipose tissue functions as a metabolically dynamic organ which regulates various biological processes including insulin sensitivity, thermogenesis, endocrine signaling, and the modulation of immune reactions [Cohen and Kajimura, 2021]. Mammals possess two types of adipose tissue, white and brown, which are distinguished by their distinct physiological roles. White adipose tissue (WAT) primarily functions as an energy reservoir, storing excess nutrients in the form of triacylglycerols (TGs) within large unilocular white adipocytes during periods of energy surplus. Under energy-deficient conditions, WAT mobilizes these lipid stores as free fatty acids to support the metabolic demands of peripheral tissues. Anatomically, WAT is distributed across various regions in the body, with major depots broadly classified into subcutaneous (SC) and visceral fat [Tchkonia *et al*., 2013]. Brown adipose tissue (BAT) represents the second major form of adipose tissue in placental mammals which is specialized for non-shivering thermogenesis. Unlike the WAT, BAT is typically localized in smaller, discrete depots such as the anterior cervical, supraclavicular, interscapular, perirenal, and perivascular regions in humans. Brown adipocytes possess unique morphological and functional features, including multilocular lipid droplets, high-density of mitochondria, and high expression of uncoupling protein 1 (UCP1). UCP1, located in the inner mitochondrial membrane, facilitates proton leak and dissipates the proton gradient as heat instead of ATP synthesis. This UCP1-dependent thermogenic mechanism, along with other futile metabolic cycles, enables BAT to expend energy and regulate systemic metabolic balance [Sakers *et al*., 2022; Cypess *et al*., 2025]. Studies utilizing positron emission tomography located BAT to several anatomical regions, including deep neck (DN), supraclavicular, axillary, paraspinal, and mediastinal depots, in adult humans [Leitner *et al*., 2017; Virtanen *et al*., 2009; Cypess *et al*., 2009].

External stimulation, such as cold exposure, physical activity, or β-adrenergic agonists, can induce WAT browning and thermogenic activation. The accumulation of cyclic adenosine monophosphate (cAMP) activates protein kinase A (PKA), which in turn phosphorylates multiple downstream targets, including key transcription factors and coactivators. This signaling cascade finally promotes lipolysis, β-oxidation, and the transcriptional activation of thermogenic genes [Ikeda *et al*., 2018].

Upon activation by adrenergic stimuli, thermogenic adipocytes consume high amounts of macro- and micronutrients to support the elevated energy demand for heat generation. We previously reported that the availability of thiamine or vitamin B1 is crucial for the efficient thermogenic activation in human primary adipocytes originating from cervical depots [Arianti *et al*., 2023] and Simpson-Golabi-Behmel syndrome (SGBS) adipocytes [Vinnai *et al*., 2025]. The inhibition of thiamine transporter (ThTr) 2 (encoded by *SLC19A3*), which is expressed exclusively in adipocytes within the human WAT and BAT, by applying its pharmacological inhibitor, fedratinib, led to reduced thermogenic capacity. However, the underlying gene expression and molecular changes of how the thermogenic activation was prevented remained unclear.

Fedratinib was developed to treat myelofibrosis because it also inhibits Janus kinase 2 (JAK2) [Blair, 2019]. Therefore, fedratinib suppresses cellular proliferation, promotes apoptosis, and inhibits the phosphorylation of downstream effectors, such as STAT3 and STAT5, in cancer cells. Pharmacokinetic studies found that fedratinib exhibits a large volume of distribution and high plasma protein binding affinity. It is primarily metabolized by cytochrome P450 enzymes, including CYP3A4 and CYP2C19, as well as flavin-containing monooxygenase 3 (FMO3), resulting in an effective half-life of 41 hours [Blair, 2019]. Despite its therapeutic potential, clinical trials of fedratinib were suspended following reports of the incidence of Wernicke’s encephalopathy, a neurological disorder linked to thiamine deficiency, which was later attributed to its off-target inhibitory effect on ThTr2 [Rodríguez-Pardo *et al*., 2015; Zhang *et al*., 2014]. In this study, we found that fedratinib altered the transcriptomic profile of human primary cervical adipocytes upon adrenergic stimulation, in which around 40 genes, including the thermogenesis-related *DIO2*, *TGM2*, and *ID1*, were downregulated. Our findings also showed that fedratinib suppressed the expression of glutamate transporter 1 (GLT1, encoded by *SLC1A2*), leading to decreased consumption and utilization of several amino acids. In addition, correlations between ThTr2 expression in SC WAT and clinical parameters highlight the importance of thiamine availability and ThTr2 function in adipocytes, affecting the metabolic fitness of the entire body.

## 2 Materials and Methods

### 2.1. Materials

All chemicals were purchased from Sigma-Aldrich (Munich, Germany) unless stated otherwise.

### 2.2. Ethics statement and obtained tissue samples

Tissue collection was approved by the Medical Research Council of Hungary (20571-2/2017/EKU) followed by the EU Member States’ Directive 2004/23/EC on presumed consent practice for tissue collection. All experiments were carried out in accordance with the guidelines of the Helsinki Declaration. Written informed consent was obtained from all participants before the surgical procedure. During thyroid surgeries, a pair of DN and SC adipose tissue samples was obtained to rule out inter-individual variations. Patients with known diabetes, malignant tumors, or with abnormal thyroid hormone levels at the time of surgery were excluded. Human adipose-derived stromal cells (hASCs) were isolated from SC and DN fat biopsies as described previously [Tóth *et al*., 2020].

### 2.3. Differentiation and treatment of hASCs

Human primary adipocytes were differentiated from the stromal vascular fraction (SVF) of adipose tissues containing hASCs according to a described protocol applying insulin, cortisol, triiodothyronine, dexamethasone, and short-term rosiglitazone (Cayman Chemicals, Ann Arbor, MI, USA cat#71740) treatment [Arianti *et al*., 2023]. In DMEM-F12-HAM medium, adipocytes were treated with a single bolus of 500 µM dibutyryl (db)-cAMP for 10 h to mimic *in vivo* cold-induced thermogenesis. Fedratinib at a 1 µM concentration was administered to inhibit ThTr2 activity.

### 2.4. RNA isolation and RNA-sequencing analysis

Cells were collected, and the total RNA was isolated as described previously [Tóth *et al*., 2020; Arianti *et al*., 2024]. The concentration and purity of the isolated RNA were checked with a Nanodrop 2000 Spectrophotometer (Thermo Fisher Scientific, Waltham, MA, USA). RNA-sequencing analysis was carried out as described in our previous study [Tóth *et al*., 2020; Arianti *et al*., 2024]. FASTQ file data were analyzed by Galaxy (https://usegalaxy.org/) [Batut *et al*., 2021]. Differentially expressed genes (DEGs) were defined based on adjusted p-values p<0.05 from DESeq2 analysis. Heatmap was generated by GraphPad 8.0 (GraphPad Software, San Diego, CA, USA) using variance stabilizing transformation (VST) score. Enrichment pathway analysis for Reactome pathways was performed by using the open platform reactome.org [Milacic *et al*., 2024]. Gene onthology (GO) and pathway enrichment analysis were also performed by using the SRplot platform (<u>SRplot - Science and Research online plot</u>) [Tang *et al*., 2023].

### 2.5. Quantitative real-time PCR (RT-qPCR)

RNA was diluted to 100 ng/µL for all samples and was reverse transcribed to cDNA by using reverse transcription kit (Thermo Fisher Scientific, 4368814) following the manufacturer’s instructions. Validated TaqMan assays used in qPCR were designed and supplied by Thermo Fisher Scientific as listed in Supplementary Table 1. qPCR was performed in Light Cycler® 480 II (Roche, Basel, Switzerland). The following conditions were set to perform the reactions: initial denaturation step at 95 °C for 1 min followed by 40 cycles of 95 °C for 12 s and 60 °C for 30 s. Gene expression values were calculated by the comparative threshold cycle (Ct) method as described in the previous publication [Arianti *et al*., 2024]. Gene expressions were normalized to the geometric mean of *ACTB* and *GAPDH*. Normalized gene expression levels equal 2^−ΔCt^.

### 2.6. Immunoblotting and densitometry

Treated cells were washed with PBS, lysed with SDS denaturing agent (50 mM Tris–HCl, 0.1% Triton X-100, 1 mM EDTA, 15 mM 2-MEA, protease inhibitors, and 5× Laemmli loading buffer). Samples were heated up to 100 °C for 10 min at a 300 rpm thermoshaker, then loaded to 10% SDS-gel for electrophoresis. Proteins were then transferred into a polyvinylidene fluoride (PVDF) membrane followed by blocking with 5% milk powder in diluted tris-buffered saline (TTBS) solution for an hour at room temperature. Next, incubation with primary antibody diluted in 0.5% milk powder in TTBS solution was done at 4 °C overnight with gentle agitation to detect the proteins of interest. The antibodies and working dilutions used in this study are listed in Supplementary Table 2. Secondary antibody solutions (horseradish peroxidase-conjugated species-corresponding secondary antibodies: goat anti-mouse IgG and goat anti-rabbit IgG) were applied in 0.5% milk powder in TTBS for an hour at room temperature. The membranes were exposed to Immobilon Western Chemiluminescent HRP Substrate (Cat. No. WBKLS0500) and the bands were detected on X-ray films (CP-BU M medical x-ray film blue, LOT 79660016) by a specific developer (DENTAMAT 500, LOT 1331) and a fixer (DENTAMAT 500, LOT 1332). For quantification of the western blot results, densitometry analysis of immunoblots was performed using the ImageJ program [Vinnai *et al*., 2023].

### 2.7. Oxygen Consumption Rate (OCR) measurement

Oxygen consumption rate (OCR) of adipocytes were measured using an XF96 oxymeter (Seahorse Biosciences, North Billerica, MA, USA) as described previously [Arianti *et al*., 2023]. After recording the baseline OCR, 500 µM db-cAMP, 1 µM fedratinib, or combination of the cAMP analogue and the inhibitor were injected to the cells. Then, stimulated OCR was recorded every 30 min for 10 h. Etomoxir (ETO) at 50 µM concentration was injected to measure ETO-resistant respiration [Nagy *et al*., 2022; Vámos *et al*., 2023]. Proton leak respiration was determined after injecting ATP synthase blocker (oligomycin) at 2 μM concentration. Cells received a single bolus of Antimycin A at 10 μM concentration for baseline correction (measuring non-mitochondrial respiration). The OCR was normalized to protein content.

### 2.8. Quantification of amino acids and calculation of their uptake by cells

The quantification of amino acids was performed following the previously described method [Arianti *et* al., 2021; Vámos *et* al., 2023]. Frozen cell culture supernatants were filtered using 3 kDa filters (Pall Corporation, Port Washington, NY, USA), and then, 10 μL of filtrate was derivatized with AccQ·Tag Ultra Derivatization Kit (Waters, Milford, MA, USA). Chromatographic separation was carried out on H-class UPLC (Waters) using AccQTag Ultra Column (2.1 × 100 mm), AccQTag Eluent A and B, and gradient provided in the AccQTag Ultra Chemistry Kit (Waters). Detection of amino acids was performed both in mass spectrometer (5500 QTRAP, Sciex) and at 260 nm in the PDA detector of the UPLC. Concentration of the amino acids was calculated with the Empower software (Waters) using a 7-point calibration curve. The flux of amino acids into or from adipocytes was calculated by comparing concentration differences measured at the starting and end point of 10 h of db-cAMP treatment with or without the presence of fedratinib.

### 2.9. Data visualization and statistical analysis

The results are expressed as mean ±SD. Normality of the distribution of the data was tested by Shapiro–Wilk test. Multiple comparisons among groups were analyzed by one-way ANOVA followed by Tukey’s *post hoc* test. The numeric data were visualized and analyzed by using GraphPad Prism 8 (GraphPad Software, San Diego, CA, USA). Graphical abstract was generated by using Biorender under the license RS28OH0HOF.

## 3 Results

### 3. 1. The expression of *SLC19A3* encoding ThTr2 was enriched in human adipocytes originating from DN and abdominal SC adipose tissues

Adipose tissue exhibits remarkable heterogeneity, consisting of various cell types including immune cells, ASCs, neuronal cells, and endothelial cells [Maniyadath *et al*., 2023]. First, we investigated the cell types that express ThTr2, encoded by *SLC19A3*, in two distinct human adipose tissue depots by utilizing publicly available data of single-cell/nuclei RNA-sequencing. We found that the expression of *SLC19A3* was exclusively enriched in the adipocytes cluster within human BAT originated from the DN depot (**Fig. 1A**) [Sun *et al*., 2020]. Consistently, analysis of *ex vivo* differentiated human brown adipocytes from eight individuals further confirmed the selective enrichment of *SLC19A3* (**Fig. 1B**) [Sun *et al*., 2020]. Next, we analyzed an open-access dataset in a recently launched platform, adiposetissue.org to investigate the distribution of *SLC19A3* expression in human WAT [Zhong *et al*., 2025]. In line with human BAT, we also found that *SLC19A3* was highly expressed in adipocytes within the abdominal WAT (**Fig. 1C**). The analysis of gene expression across various human tissues showed a strong specificity of *SLC19A3* expression in adipocytes marked by high rank of its expression level (**Fig. 1D**). These data suggest that *SLC19A3* exhibits adipocyte-specific expression across both BAT and WAT depots, highlighting its potential role as a key molecular component in adipocyte function and metabolic regulation.

**Figure 1.**
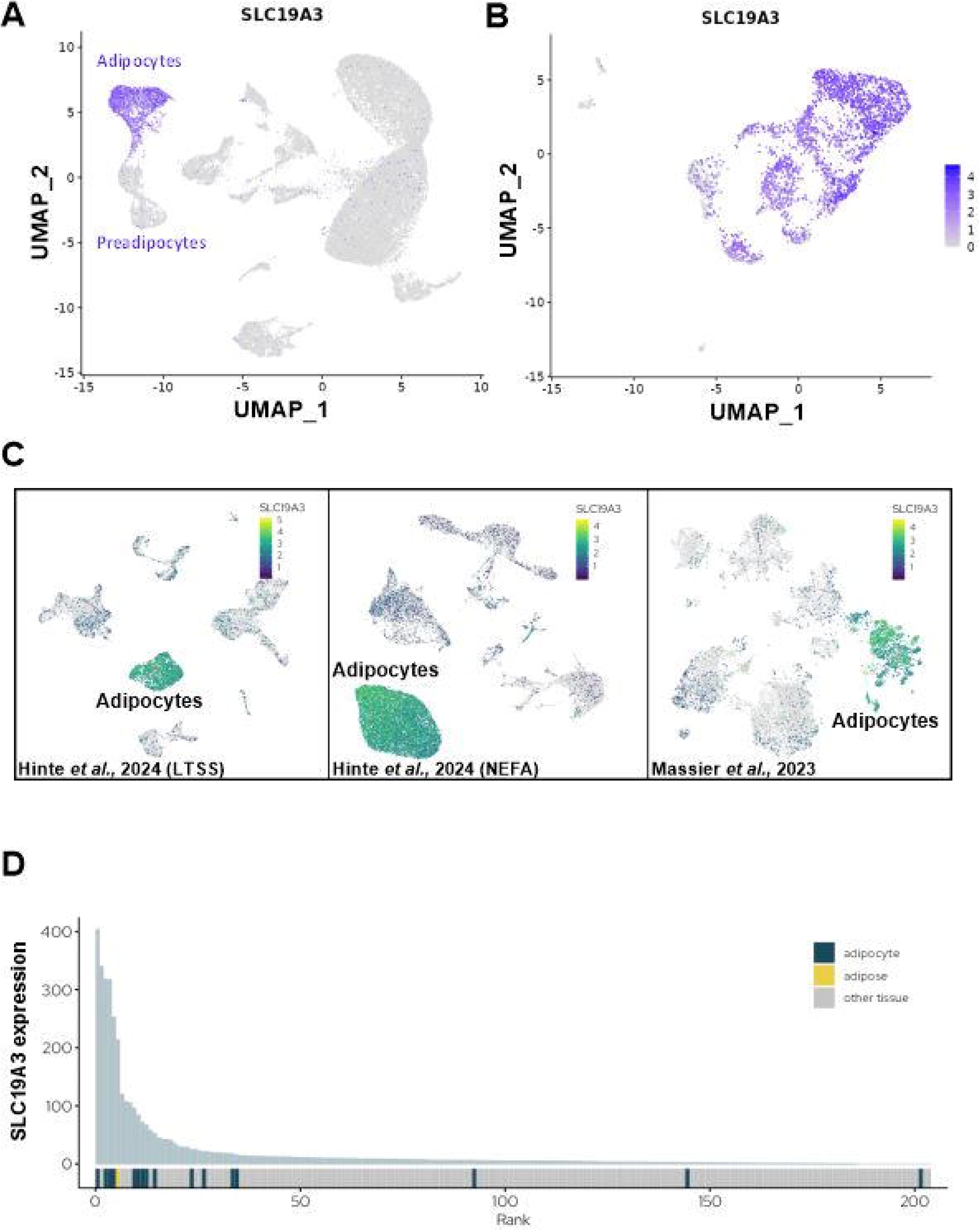
Tissue-specific expression of Thiamine Transporter (ThTr) 2 (encoded by *SLC19A3*) in human adipose depots. (A-B) Embedding plots showing the expression of *SLC19A3* by single-nuclei RNA-sequencing from human brown adipose tissue (A) and differentiated brown adipocytes (B) (data was retrieved from *batnetwork.org*) [Sun *et al*., 2020]. (C) Embedding plots of single-cell expression profile of *SLC19A3* in human abdominal subcutaneous white adipose tissue (data was retrieved from *adiposetissue.org*) [Zhong *et al*., 2025; Hinte *et al.,* 2024*; Massier et al*., 2023], indicating major cell lineages and the expression level of *SLC19A3* in these lineages. (LTSS: Leipzig two-step surgery; NEFA Trial NCT01727245). (D) Tissue specificity graph presenting data from the FANTOM consortium, highlighting the expression levels of genes in adipose tissue relative to other human organs and tissues, emphasizing which genes are actively expressed in the specific tissues (data was retrieved from *adiposetissue.org*) [Zhong *et al*., 2025].

### 3.2. Fedratinib prevented the elevation of thermogenic markers expression and oxygen consumption in human cervical-derived adipocytes during adrenergic stimulation

The specific enrichment of *SLC19A3* in adipocytes cluster prompted us to further analyze its importance in the gene expression regulation of thermogenic adipocytes. We studied the effect of pharmacological inhibition of ThTr2 on the expression of thermogenic markers by applying a potent inhibitor, fedratinib [Arianti *et al*., 2023; Zhang *et al*., 2014]. We found that the mRNA and protein expression of UCP1, PGC1a, and DIO2 were hampered by fedratinib during adrenergic stimulation in both SC and DN-derived adipocytes (**Fig. 2A, B**). Next, we analyzed the effect of fedratinib on cellular respiration and found that the db-cAMP-stimulated elevation of maximal respiration tended to be abrogated by fedratinib. More importantly, proton leak respiration reflecting UCP1-dependent heat generation was significantly hampered by fedratinib during db-cAMP-driven activation in both SC and DN-derived adipocytes (**Fig. 2C**), suggesting that ThTr2 plays a crucial role in sustaining the thermogenic capacity of human neck-derived adipocytes during adrenergic stimulation.

**Figure 2.**
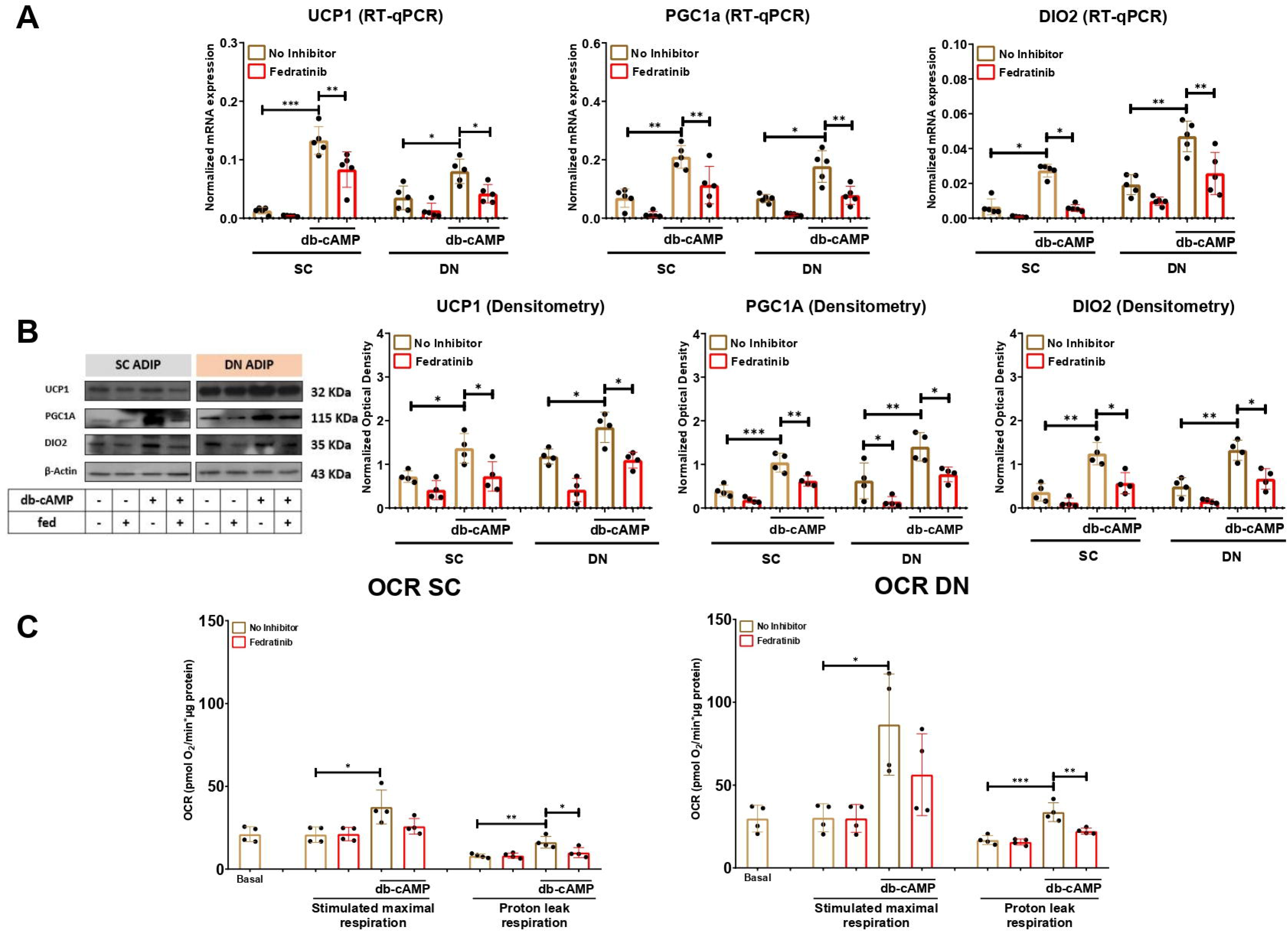
The effect of fedratinib (fed) on the expression of thermogenic markers in subcutaneous (SC) and deep neck (DN)-derived adipocytes after 10 h of dibutyryl (db)-cAMP-induced thermogenesis. (A) The mRNA expression of *UCP1, PGC1a,* and *DIO2* in SC and DN-derived adipocytes upon 10 h of thermogenic activation induced by db-cAMP was analyzed by RT-qPCR. (B) The protein expression of UCP1, PGC1a, and DIO2 was assessed by immunoblotting and quantified by optical densitometry. (C) Oxygen consumption rate (OCR) was measured by Seahorse extracellular flux analysis at basal, maximal db-cAMP-driven stimulation, and the proton leak respiration after oligomycin addition in both SC and DN-derived adipocytes. n=5 for RT-qPCR, n=4 for immunoblotting, and n=4 for OCR. The original pictures of the full-length blots are displayed in Supplementary Fig. 4. Statistical analysis was performed by one-way ANOVA followed by Tukey’s *post hoc* test, *p<0.05, **p<0.01, ***p<0.001.

### 3.3. Fedratinib alters the gene expression profile of human neck-derived adipocytes during adrenergic stimulation

Next, we aimed to investigate the impact of fedratinib on the gene expression profile of human neck-derived adipocytes during thermogenic activation by performing bulk RNA-sequencing analysis. Our data showed that among 758 genes which were upregulated by db-cAMP in SC adipocytes [Arianti *et al*., 2024], 14 genes were suppressed by fedratinib (**Fig. 3A, left panel**). In DN-derived adipocytes, there were 927 genes that were upregulated by db-cAMP [Arianti *et al*., 2024], and 9 of those were hampered by fedratinib (**Fig. 3A, right panel**). Our data also showed that the expression of 26 and 32 genes were not influenced by db-cAMP, but they were suppressed by fedratinib in SC and DN-derived adipocytes, respectively (**Fig. 3A**). We found 15 and 7 genes that were upregulated by fedratinib in SC and DN-derived adipocytes, respectively (**Supplementary Table 3 and 4**).

**Figure 3.**
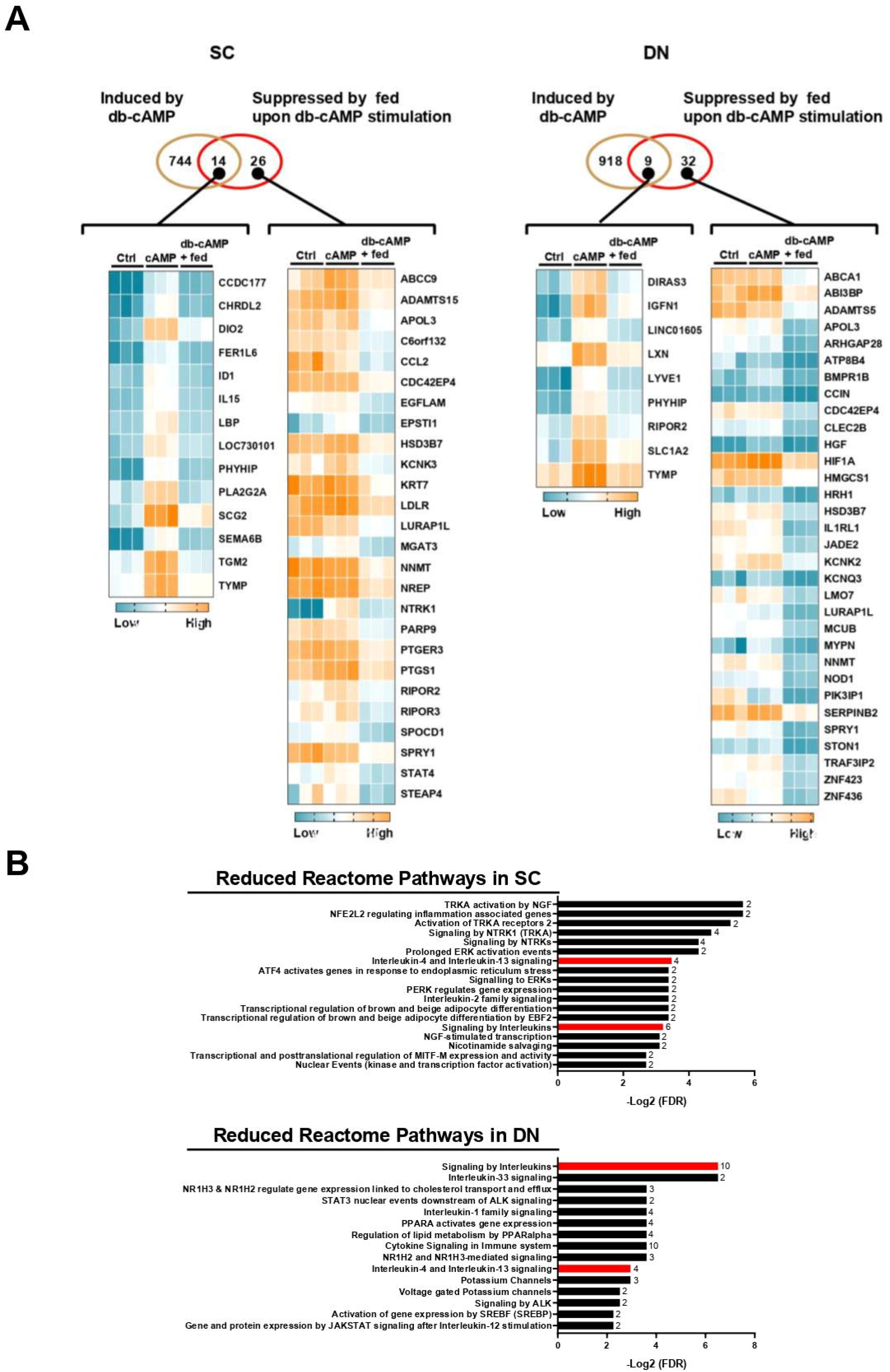
Effect of fedratinib (fed) on the global gene expression profile of subcutaneous (SC) and deep neck (DN)-derived adipocytes after 10 h of thermogenic activation induced by dibutyryl (db)-cAMP. (A) Venn diagrams showing the numbers of upregulated genes by db-cAMP (brown circle) or downregulated genes by fed (red circle) in SC or DN-derived adipocytes, visualized by the heat map displaying the differentially expressed genes (DEGs) in both SC (left panel) or DN (right panel) adipocytes. (B) Reactome pathways in which DEGs that were suppressed by fed during db-cAMP-stimulated thermogenesis were enriched in SC (upper panel) or DN (lower panel)-derived adipocytes (The common pathways are colored in red; numbers next to columns indicate the number of DEGs enriched in the pathway). FDR: false discovery rate.

Furthermore, to gain insight into the broader biological effect of fedratinib treatment, we performed Reactome pathway enrichment analysis on the identified DEGs which were suppressed by fedratinib. We found various overrepresented biological pathways, including transcriptional regulation of brown and beige adipocyte differentiation by EBF2, signaling to ERKs, activation of TRKA receptors, and interleukin-2 family signaling in SC (**Fig. 3B, top panel**), and activation of gene expression by SREBF (SREBP), PPARA activates gene expression, and voltage-gated potassium channels in DN-derived adipocytes (**Fig. 3B, bottom panel**). We found three affected pathways which were commonly enriched in SC and DN-derived adipocytes including signaling by interleukins, interleukin-4 and interleukin-13 signaling, and nicotinamide salvaging pathways (**Fig. 3B**). We also performed GO and pathway enrichment analysis and found that fedratinib suppressed genes which were involved in several biological processes (positive regulation on external stimulus and NAD biosynthesis via nicotinamide riboside salvage pathway), molecular function (ATPase dependent transmembrane transport complex), cellular component, receptor ligand activity, and pathways (regulation of lipolysis in adipocytes and JAK-STAT signaling pathway) in SC adipocytes (**Supplementary Fig. 1**). In DN adipocytes, genes which were enriched in biological processes of regulation of GTP binding and positive regulation of carbohydrate metabolic process, the molecular function of glycosaminoglycan binding, cellular component of astrocyte projection, and the pathway of primary bile acid biosynthesis were suppressed by fedratinib (**Supplementary Fig. 2**). These findings suggest that fedratinib regulates thermogenic activation in human neck-derived adipocytes by suppressing gene expression and signaling pathways involved in brown adipocyte differentiation, immune signaling, and metabolic regulation.

### 3.4. Fedratinib hampered the expression of recently identified thermogenesis-related genes during adrenergic stimulation

Our RNA-sequencing analysis showed that the expression of newly identified thermogenesis-related genes, including transglutaminase 2 (*TGM2*) [Mádi *et* al., 2017; Lénart *et al*., 2022; Svensson *et al*., 2011] and inhibitor of DNA binding 1 (*ID1*) [Arianti *et* al., 2024], was abrogated by fedratinib during adrenergic stimulation. Next, we validated that the expression of TGM2 and ID1 was significantly increased by db-cAMP; however, their upregulation was prevented by fedratinib during db-cAMP-driven activation at both mRNA (**Fig. 4A**) and protein levels (**Fig. 4B**) in both SC and DN-derived adipocytes. These findings suggest that the decreased expression of TGM2 and ID1 contribute to the thermogenesis inhibiting effect of fedratinib in human neck-derived adipocytes in response to adrenergic stimulation. Interestingly, our RNA-sequencing data showed that the expression of nicotinamide N-methyltransferase (NNMT), which is involved in nicotinamide salvaging pathway, was decreased by fedratinib in both SC and DN-derived adipocytes which was confirmed by RT-qPCR and western blot (**Supplementary Fig. 3**).

**Figure 4.**
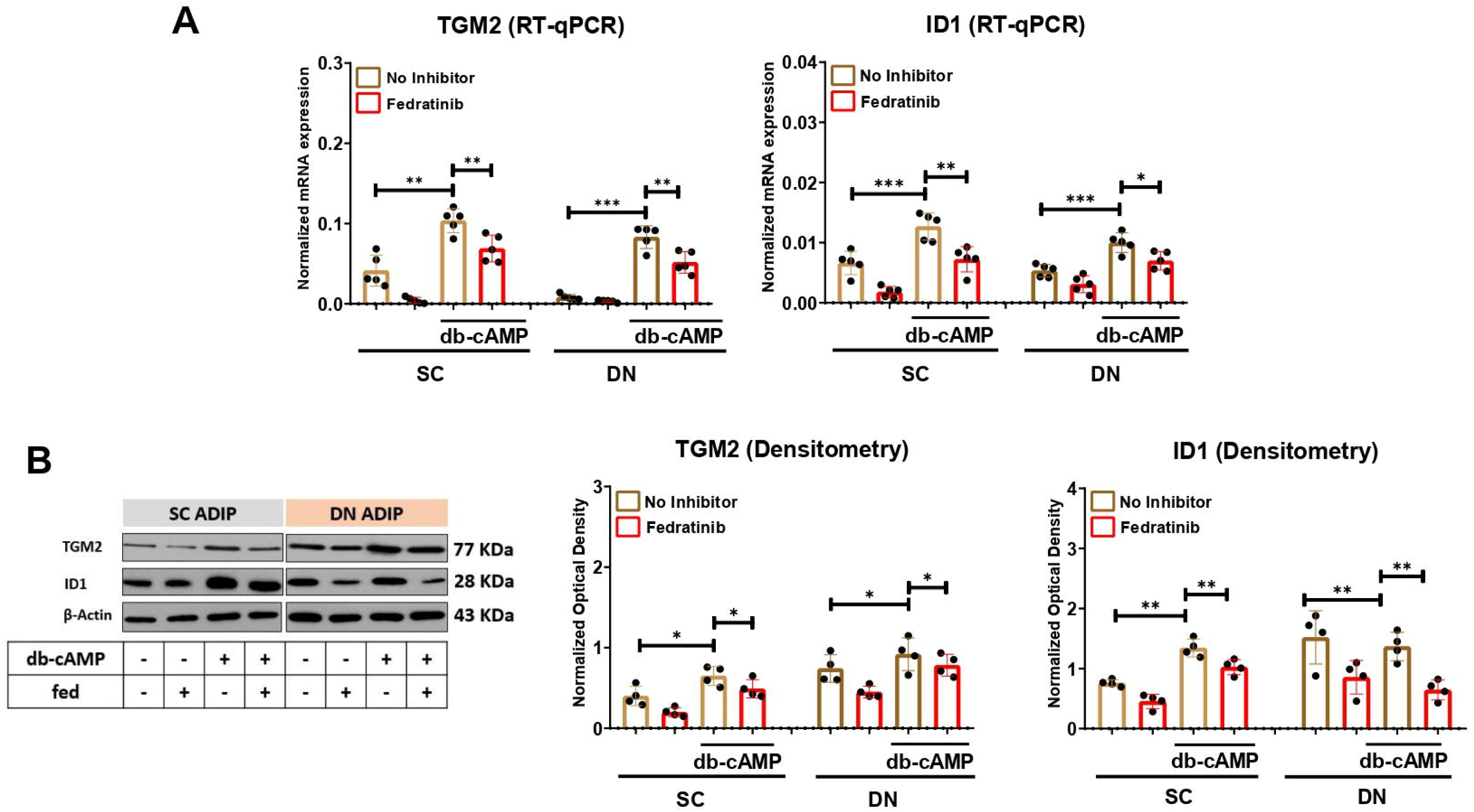
The effect of fedratinib (fed) on the expression of thermogenesis-related genes in subcutaneous (SC) and deep neck (DN)-derived adipocytes after 10 h of thermogenic activation induced by dibutyryl (db)-cAMP. (A) The mRNA expression of *TGM2,* and *ID1* was analyzed by RT-qPCR. (B) Protein expression of TGM2 and ID1 was detected by immunoblotting. n=5 for RT-qPCR, and n=4 for immunoblotting. The original pictures of the full-length blots are displayed in Supplementary Fig. 5. Statistical analysis was performed by one-way ANOVA followed by Tukey’s *post hoc* test, *p<0.05, **p<0.01, ***p<0.001.

### 3.5. Amino acid transport and utilization were inhibited by fedratinib during adrenergic stimulation

Our RNA-sequencing data showed that the expression of solute carrier family 1 member 2 (*SLC1A2*), which encodes glutamate transporter (GLT) 1, was decreased by fedratinib (**Fig. 3A**). Next, we investigated the amino acid consumption by the human neck-derived adipocytes upon adrenergic stimulation. We found that the consumption of glutamine (Gln), arginine (Arg), glycine (Gly), cysteine (Cys), tyrosine (Tyr), and valine (Val) was decreased during adrenergic stimulation in the presence of fedratinib with more pronounced effect observed in DN-derived adipocytes (**Fig. 5A**). In parallel, the etomoxir (ETO)-resistant respiration reflecting glucose and amino acids utilization was also reduced by fedratinib during db-cAMP-driven activation in DN adipocytes (**Fig. 5B**). We also validated our RNA-sequencing data showing that the mRNA and protein expression of GLT1 was abrogated by fedratinib upon adrenergic stimulation (**Fig. 5C, D**). The L-type amino acid transporter (LAT) family is a major entry gate for amino acids through the cell membrane, which consists of 4 members: LAT1-4 [Zhao *et al*., 2023]. LAT1 and 2 are members of the SLC7 family, encoded by *SLC7A5* and *SLC7A8*, respectively, while LAT3 and 4 are classified to the SLC43 family, encoded by *SLC43A1* and *SLC43A2*, respectively [Bodoy *et al*., 2005; Babu *et al*., 2003]. Adrenergic stimulation by db-cAMP significantly increased the expression of LAT1 and 2 in both SC and DN-derived adipocytes; however, only LAT2 expression was affected by fedratinib (**Fig. 5E**). The expression of LAT3 and 4 was not affected by either db-cAMP or fedratinib (**Fig. 5E**). Next, we validated our RNA-sequencing data showing that the mRNA and protein levels of LAT2 were significantly hampered by fedratinib upon adrenergic stimulation (**Fig. 5F, G**).

**Figure 5.**
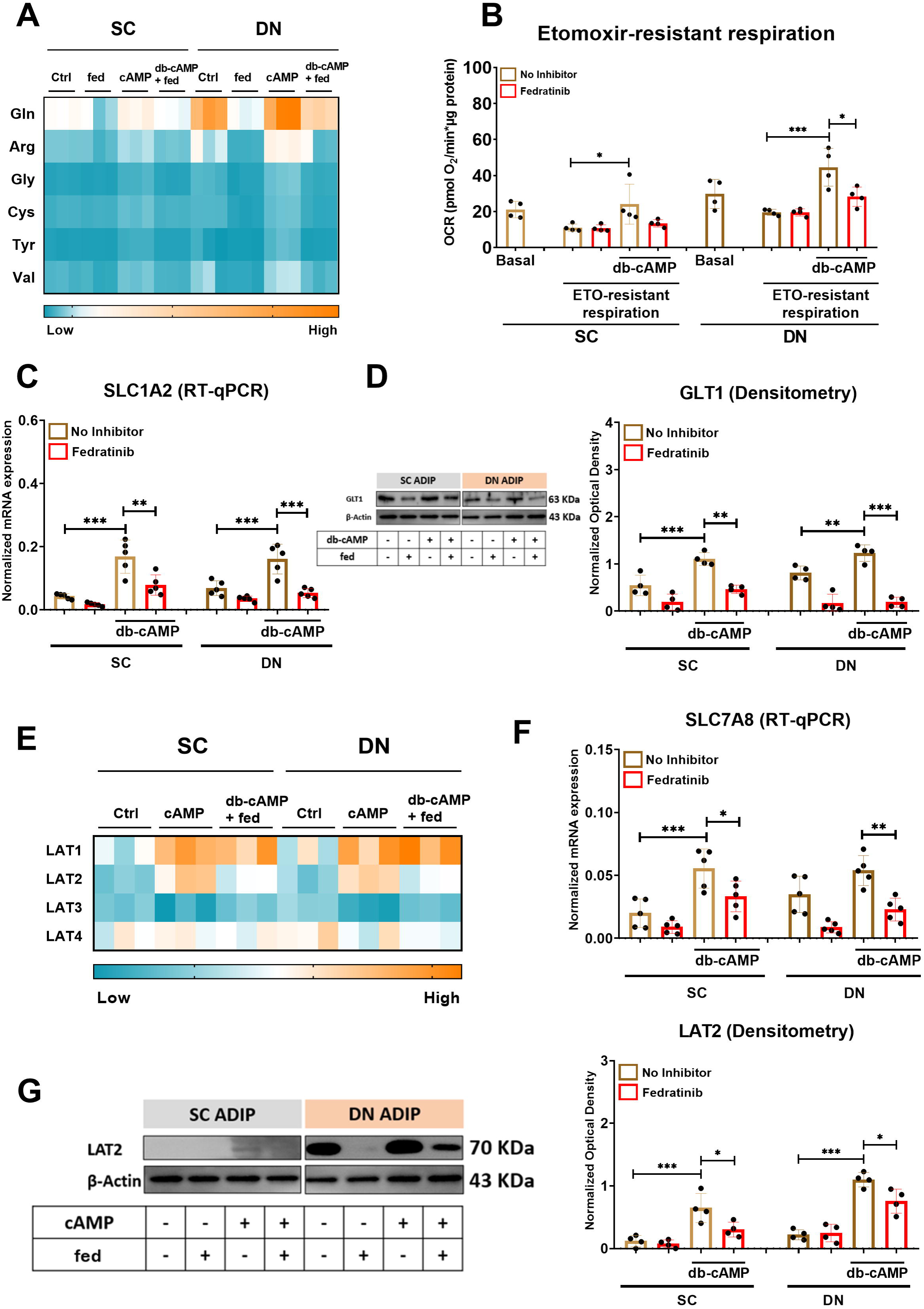
Effect of fedratinib (fed) on the expression of amino acid transporters and utilization of amino acids in subcutaneous (SC) and deep neck (DN)-derived adipocytes during thermogenic activation by dibutyryl (db)-cAMP. (A) Heat map displaying the amino acid consumption. (B) Etomoxir (ETO)-resistant respiration was measured by Seahorse extracellular flux analysis, n=4. (C) mRNA expression of glutamate transporter 1 (GLT1) (encoded by *SLC1A2*) was analyzed by RT-qPCR, n=5. (D) GLT1 protein expression was assessed by immunoblotting, n=4. (E) Heat map displaying the mRNA expression of L-amino acid transporters (LAT) 1-4 in SC and DN-derived adipocytes based on the RNA-seq data analyzed by DESeq2. (F) The mRNA level of the LAT2 (encoded by *SLC7A8*) was quantified by RT-qPCR, n=5. (G) The protein expression of LAT2 was analyzed by immunoblotting, n=4. The original pictures of the full-length blots are displayed in Supplementary Fig. 6. Statistical analysis by one-way ANOVA followed by Tukey’s *post hoc* test, *p<0.05, **p<0.01, ***p<0.001.

### 3.6. The expression of ThTr2 in human WAT positively associates with metabolic health

To evaluate the clinical significance of ThTr2, we analyzed data available at adiposetissue.org, which includes large-scale transcriptomic and proteomic datasets from human abdominal SC WAT [Zhong et al., 2025]. The association analyses were organized into three major clinical categories: (I) anthropometric traits, such as body mass index (BMI) and waist-hip ratio (WHR); (II) circulating diagnostic markers, including inflammatory cytokines and metabolic hormones (e.g., insulin); and (III) tissue-specific expression patterns, focused on genes and proteins enriched in abdominal WAT (**Fig. 6A**). We observed that ThTr2 expression at both the mRNA and protein levels showed inverse correlations with body composition measures, such as BMI and WHR, indicating that ThTr2 is more highly expressed in individuals with lower adiposity. Furthermore, *SLC19A3* expression inversely correlated with Homeostatic Model Assessment for Insulin Resistance (HOMA-IR), pro-inflammatory markers (e.g., CRP), circulating low-density lipoprotein (LDL), insulin, glucose, TG, cholesterol, and HbA1c, but positively correlated with beneficial metabolic markers, such as high-density lipoprotein (HDL) cholesterol, suggesting a link between higher ThTr2 expression and a healthier systemic metabolic profile (**Fig. 6A, left and middle panels**). In addition, ThTr2 expression was negatively correlated with fat cell volume but positively associated with lipolysis, indicating that enriched expression of ThTr2 is beneficial to prevent the hypertrophy of WAT (**Fig. 6A, right panel**). In parallel, a meta-analysis comparing ThTr2 expression in WAT between non-obese individuals (BMI <30 kg/m²) and patients with obesity (BMI ≥30 kg/m²) revealed higher levels of ThTr2 in non-obese individuals (**Fig. 6B**). Lastly, in four longitudinal weight loss intervention cohorts, including two diet-induced and two surgery-induced models, we found that the expression of ThTr2 was elevated upon bariatric surgery or 8 weeks after diet (**Fig. 6C**), further reinforcing the link between improved metabolic state and the upregulation of ThTr2. These findings in clinical parameters suggest that ThTr2, which is specifically enriched in the adipocyte cluster within WAT and BAT, is a potential molecular marker of metabolic and adipose tissue health.

**Figure 6.**
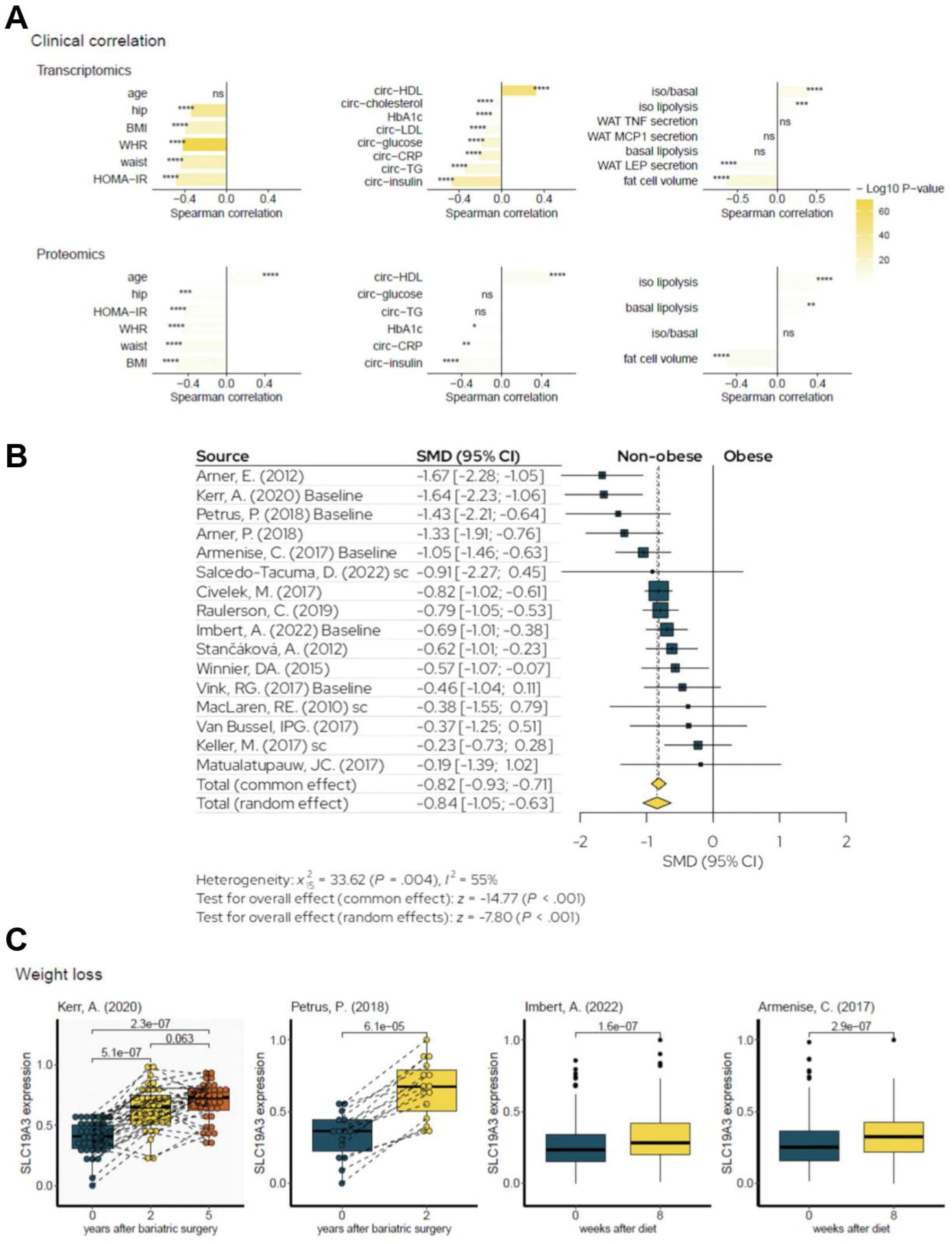
Correlation of Thiamine transporter 2 (ThTr2 encoded by *SLC19A3*) expression in abdominal subcutaneous white adipose tissue (WAT) with clinical parameters. Data was retrieved from *adiposetissue.org* [Zhong *et al*., 2025]. (A) The transcriptome (upper panels) and proteome (bottom panels) analyses were categorized into three groups: anthropometric (left panels), focusing on *SLC19A3* gene and protein expression in correlation with measurements of body distribution parameters (e.g., BMI); circulating diagnostic markers (middle panels), examining molecules present in the bloodstream (e.g., hormones); and tissue-specific responses (right panels), highlighting genes and proteins predominantly expressed in abdominal WAT. This classification helps to differentiate systemic metabolic regulation from localized tissue functions and their associations with body composition. Data in upper and bottom right panels were obtained from abdominal WAT-derived adipocytes differentiated *ex vivo*. BMI: body mass index; HOMA-IR: Homeostatic Model Assessment for Insulin Resistance, WHR: waist-hip ratio; HDL: high-density lipoprotein; HbA1c: hemoglobin A1c; LDL: low-density lipoprotein; TG: triglyceride TNF: tumor necrosis factor; MCP1: monocyte chemoattractant protein 1; LEP: leptin; CRP: C-reactive protein; iso: isoproterenol. (B) Meta-analysis forest plot comparing the of ThTr2 mRNA levels in people living with or without obesity based on the respective BMI cutoffs (<30 kg/m^2^ vs. ≥30 kg/m^2^). SMD: Standardized mean differences [Data referenced from Arner *et al.,* 2012; Kerr *et al.,* 2020; Petrus *et al.,* 2018; Arner *et al.,* 2018; Armenise *et al.,* 2017; Salcedo-Tacuma *et al.,* 2022; Civelek *et al.,* 2017; Raulerson *et al.,* 2019; Imbert *et al.,* 2022; Stančáková *et al.,* 2012; Winnier *et al.,* 2015; Vink *et al.,* 2017; MacLaren *et al.,* 2010; Van Bussel *et al.,* 2017; Keller *et al.,* 2017; and Matulalupauw *et al.,* 2017]. (C) Boxplots for 2 surgery-induced and 2 diet-induced weight loss cohorts indicate that *SLC19A3* mRNA expression in WAT was increased after the interventions (Data referenced from [Kerr *et al*., 2020; Petrus *et al*., 2018; Imbert *et al*., 2022; Armenise *et al*., 2017] (*adiposetissue.org*).

## 4 Discussion

Thermogenically active brown adipocytes are very energy-spending cells, utilizing readily available metabolic substrates to support heat generation [Townsend and Tseng, 2014]. Adrenergic stimulation, such as induced by cold exposure, increases the consumption of macronutrients such as glucose [Orava *et al*., 2011], fatty acids [Wu *et al*., 2006; Bartelt *et al*., 2011], and amino acids [Arianti *et al*., 2021; Jersin *et* al., 2021; Yoneshiro *et al*., 2019; Park *et al*., 2023] and micronutrients such as thiamine [Arianti *et al*., 2023; Vinnai *et al*., 2025]. Active brown and beige adipocytes require increased levels of thiamine to enhance the activity of thiamine pyrophosphate (TPP)-dependent enzymes, pyruvate and α-ketoglutarate dehydrogenases, which are critical for generating reduced equivalents that drive the generation of mitochondrial proton gradient. In addition, a critical step in the breakdown of branched-chain amino acids, which are crucial nutrients during thermogenesis, is catalyzed by the TPP-requiring branched-chain alpha-ketoacid dehydrogenase (BCKDH) [Verkerke *et al*., 2024]. Thiamine enters the cells mostly via cell membrane transporters of the SLC19 family [Manzetti *et al*., 2014]. Our previous study highlighted that ThTrs were highly expressed in thermogenic adipocytes [Tóth *et al*., 2020]. An RNA-seq-based screening identified ThTr2 as specifically enriched in human abdominal WAT, where its high expression levels strongly correlate with mitochondrial gene expression, suggesting a potential association between the abundance of ThTr2 and brown/beige adipocyte function [Pereira *et al*., 2021]. Furthermore, single-cell RNA-sequencing analysis (**Fig. 1**) showed that ThTr2 was exclusively enriched in the adipocytes cluster in both human BAT and WAT suggesting its importance in facilitating rapid responses to thermogenic stimuli that demand increased energy expenditure.

Fedratinib was previously used in the treatment of myelofibrosis as a JAK2 inhibitor [Blair, 2019]; however, its use was suspended following reported incidences of Wernicke’s encephalopathy, a severe neurological disorder associated with thiamine deficiency [Zhang *et al*., 2014]. This raises the possibility that the effects of fedratinib on thermogenic activation may result from the inhibition of an activated tyrosine kinase, particularly given that JAK2^−/−^ mice fail to upregulate Ucp1 in BAT following high-fat diet feeding or cold exposure [Shi *et al*., 2016]. Our RNA-seq analysis showed that the expression of *STAT4* was decreased in response to fedratinib when we compared db-cAMP and the combination of db-cAMP and fedratinib treatments (**Fig. 3a**). Reactome pathway analysis also highlighted that genes involved in gene and protein expression by JAK/STAT signaling after interleukin-12 stimulation were significantly affected by fedratinib (**Fig. 3b**). However, the possibility that fedratinib impairs thermogenesis at least partially because of limited thiamine uptake is supported by previous and current findings. On one hand, we showed that fedratinib decreased facilitated thiamine influx during adrenergic stimulation in human neck-derived adipocytes [Arianti *et al*. 2023]. In line with our findings, Zhang *et al*. (2014) also reported that fedratinib significantly inhibited thiamine uptake in CACO2 cells. Taken together, these findings suggest that fedratinib may impair thermogenic activation through both JAK2-dependent signaling pathways and inhibition of thiamine uptake, highlighting the need to consider both mechanisms in interpreting its metabolic effects.

In our previous study, we demonstrated that thiamine availability, transported primarily via ThTr1 and ThTr2, is essential for a robust thermogenic response following db-cAMP stimulation in human primary adipocytes derived from SC and DN fat depots [Arianti *et al*., 2023]. Pharmacological inhibition of ThTrs resulted in decreased expression of key thermogenic genes, including *UCP1*, *PGC1*α, and *DIO2* as well as reduction in UCP1-dependent heat production as measured by proton leak respiration. Additionally, we also confirmed that sufficient thiamine availability is crucial for the effective thermogenic activation in SGBS adipocytes [Vinnai *et al*., 2025]. In line with our findings, a recent study identified thiamine, along with pantothenic acid and riboflavin, as critical vitamins supporting the adipogenic differentiation of human dermal fibroblasts and ASCs into brown-like adipocytes [Vinnai *et* al., 2023; Takeda and Dai, 2024]. The current study reveals the transcriptomic alteration by fedratinib that may underlie the reduced thermogenic capacity in human neck-derived adipocytes. The expression of thermogenesis-related genes such as *TGM2* [Lénart *et al*., 2022], *ID1* [Arianti *et al*., 2024], *KCNK3* [Shinoda *et al*., 2015], and *HMGCS1* [Balaz *et al*., 2019] was suppressed by fedratinib. Transcriptomic analysis showed that the expression of *TGM2* (encoding transglutaminase 2) was higher in human BAT originating from DN as compared to SC WAT of neck-origin [Svensson *et al*., 2011]. Lénart *et al*. (2022) reported that TGM2^−/−^ mice showed a lower cold exposure tolerance, enlarged adipocyte size compared to wild-type mice, and decreased mRNA expression of the key brown adipocyte marker Ucp1. The db-cAMP-stimulated upregulation of ID1 and 3 was shown to be critical for thermogenic activation in human neck-derived adipocytes [Arianti *et al*., 2024]. Expression of KCNK3, which encodes a pH-dependent, voltage-insensitive potassium channel protein, was significantly higher in human brown as compared to white adipocytes [Shinoda *et al*., 2015] and was induced in the supraclavicular BAT depots from six subjects under prolonged cold exposure (19 °C) versus thermoneutral conditions (30 °C) [Chondronikola *et al*., 2014]. Balaz *et al*. found that the knockdown of HMGCS1 in both mice and human brown adipocytes led to the reduced UCP1 expression and mitochondrial respiration. In addition to the known thermogenesis-related genes, our RNA-sequencing analysis showed that fedratinib decreased the expression of *NNMT* in human DN-derived adipocytes (**Fig. 3 and Supplementary Fig. 3**). *NNMT* encodes nicotinamide N-methyltransferase, a key enzyme that catalyzes the methylation of nicotinamide to form N-methyl nicotinamide, by using adenosylmethionine as methyl donor. Previous studies reported that NNMT knockdown protected against diet-induced obesity in mice [Kraus *et al*., 2014], and its expression is inversely correlated with thermogenic markers such as UCP1 and PGC1a [Jia *et al*., 2022]. The TPP requiring transketolase is a key enzyme in the non-oxidative branch of the pentose phosphate pathway that provides ribose 5-phosphate for nucleotide, such as NAD^+^ synthesis

[Alexander-Kaufman and Harper, 2009]. We speculate that the fedratinib-induced downregulation of NNMT is compensatory to preserve sufficient levels of intracellular NAD which is critical to maintain vital metabolic reactions.

Our data also demonstrated that fedratinib decreased amino acids consumption and utilization upon adrenergic stimulation, likely through the downregulation of GLT1 and LAT2. GLT1/SLC1A2 is widely expressed in the brain and our current data showed that it is also expressed in human adipocytes. The role of GLT1 in the clearance of the excitatory neurotransmitter glutamate from the synaptic clefts in the central nervous system has been well-elucidated [Rimmele and Rosenberg, 2016], however, the importance of this transporter function in adipocytes remains underexplored. Notably, a previous study reported that thiamine deficiency leads to the loss of astrocytic GLT1, a primary feature of Wernicke’s encephalopathy [Jhala *et al*., 2014]. In line with these findings, our results show that fedratinib, which is a potent inhibitor of ThTr2, also reduces GLT1 expression in adipocytes, suggesting a potential mechanistic link between thiamine availability and GLT1 regulation in peripheral tissues.

The expression of ThTr2 in human abdominal SC WAT correlates with metabolic health marked by negative correlation with BMI, WHR, HOMA-IR, LDL, and cholesterol (**Fig. 6**) [Zhong *et al*., 2025]. Interestingly, the expression of ThTr2 is also inversely correlated with WAT leptin secretion, suggesting a potential role in modulating leptin resistance. Data from four cohorts also demonstrates that ThTr2 expression is elevated after bariatric surgery and weight-loss-initiating dietary interventions. This finding is in accordance with previous studies reporting that individuals with obesity frequently exhibit marked thiamine deficiency prior to bariatric surgery [Nath *et al*., 2017; Flancbaum *et al*., 2006; Peterson *et al*., 2016]. A recent publication reported that fecal thiamine levels exhibited a positive correlation with HDL-C and a negative correlation with BMI, HOMA-IR, fasting serum insulin, TG, and propionic acid levels [Xia *et al*., 2025].

Our findings highlighted that fedratinib altered the transcriptomic profile of human neck-derived adipocytes during adrenergic stimulation, potentially by inhibiting ThTr2. Although our findings provide potentially useful molecular and clinical insights, we cannot rule out that fedratinib may also have currently unknown off-target effects in adipocytes. Therefore, to more precisely reveal the transcriptional regulatory role of ThTr2-mediated thiamine influx in thermogenesis, particularly in human adipocytes, future studies utilizing targeted approaches, such as siRNA-mediated knockdown or CRISPR-Cas9 gene editing, are required. Future investigations will be instrumental to advance our understanding regarding the role of ThTr2 in adipose tissue biology and to open a better therapeutic strategy by targeting this transporter.

## 5 Conflict of Interest

The authors declare that the research was conducted in the absence of any commercial or financial relationships that could be construed as a potential conflict of interest.

## 6 Author Contributions

G. K — conceptualization, methodology, investigation, data curation, validation, visualization, writing - original draft; E. Ko. — methodology, investigation, writing - original draft; R. Al. — methodology, writing - original draft; M. S. — methodology, writing - original draft; F. G. — methodology, resources, writing – review and editing; R. C. — investigation, writing - original draft; É. C. — methodology, investigation, funding acquisition, writing – review and editing; S. P. — methodology, investigation, writing – review and editing; L. F. — conceptualization, supervision, writing – review and editing; R. Ar. — conceptualization, methodology, investigation, funding acquisition, visualization, writing – review and editing; E. Kr. — conceptualization, project administration, funding acquisition, supervision, writing – review and editing.

## 7 Funding

This research was conducted with the support of the National Research, Development and Innovation Office (NKFIH-FK145866, FK134605, and PD146202) of Hungary, the János Bolyai Research Scholarship of the Hungarian Academy of Sciences, the University of Debrecen Program for Scientific Publication, and the National Academy of Scientist Education Program of the National Biomedical Foundation under the sponsorship of the Hungarian Ministry of Culture and Innovation. S.P. was supported by the project TKP2021-NKTA-34, which has been implemented with the support provided by the Ministry of Culture and Innovation of Hungary from the National Research, Development and Innovation Fund, financed under the TKP2021-NKTA funding scheme.

## 8 Acknowledgments

We thank Dr. Zsolt Sarang for his exceptional help in reviewing the manuscript before its submission and Jennifer Nagy for technical assistance.

## 10 Supplementary Material

Supplementary Material should be uploaded separately on submission, if there are Supplementary Figures, please include the caption in the same file as the figure. Supplementary Material templates can be found in the Frontiers Word Templates file.

Please see the Supplementary Material section of the Author guidelines for details on the different file types accepted.

## 12 Data Availability Statement

The RNA-seq datasets generated and analyzed for this study can be found in the Sequence Read Archive (SRA) database [https://www.ncbi.nlm.nih.gov/sra] under accession number PRJNA1093362.

## Notes

### Competing Interest Statement

The authors have declared no competing interest.

