## Supplementary Material for "Thiamine transporter 2 and Janus kinase 2 inhibitor, fedratinib suppresses thermogenic activation of human neck area-derived adipocytes"


### 1 Supplementary Figures and Tables

#### 1.1 Supplementary Figures

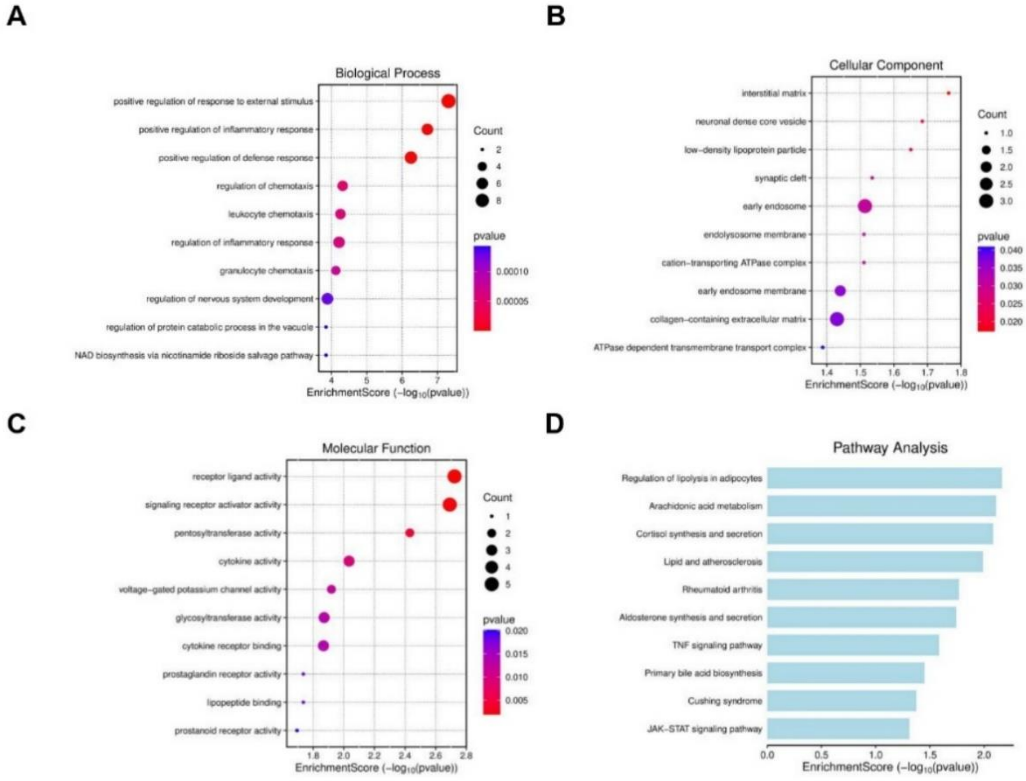


**Supplementary Figure 1. Gene Ontology enrichment in subcutaneous (SC)-derived adipocytes** (A) Biological processes, (B) cellular components, (C) molecular functions, and (D) pathway analysis of the differentially expressed genes that were suppressed by fedratinib during dibutyryl-cAMP-stimulated thermogenesis, enriched in SC-derived adipocytes [Data cited from www.bioinformatics.com.cn/SRplot].

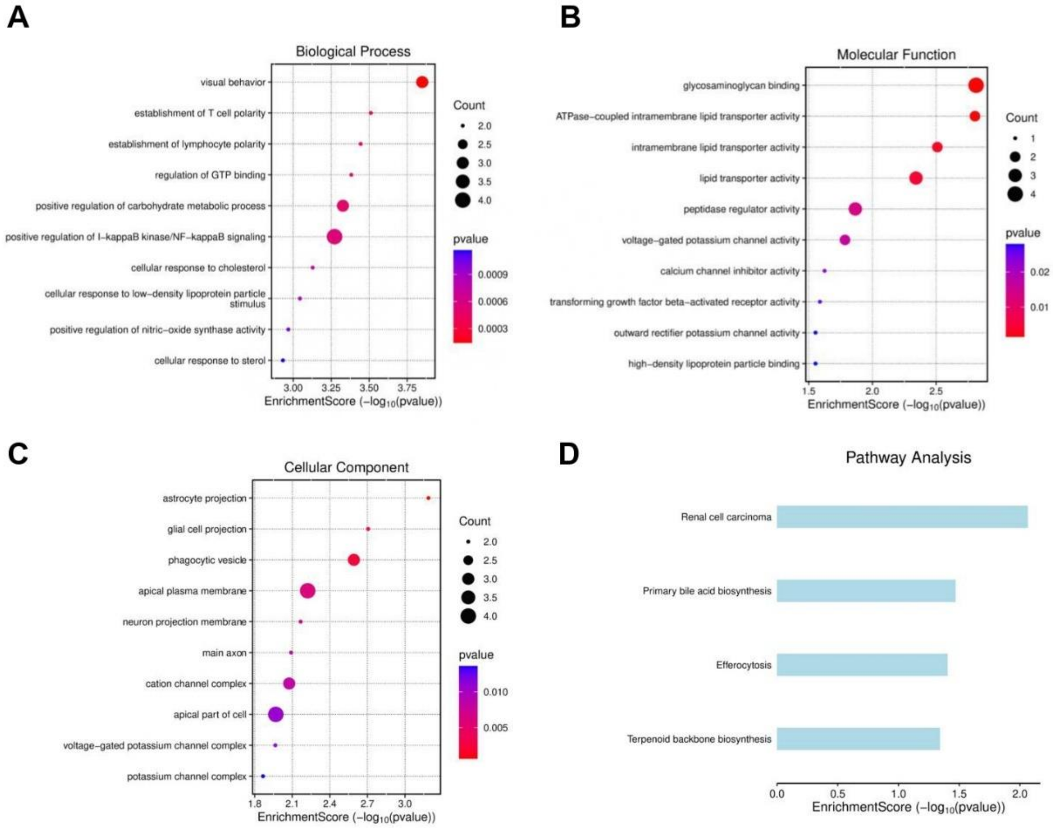


**Supplementary Figure 2. Gene Ontology enrichment in deep neck (DN)-derived adipocytes** (A) Biological processes, (B) cellular components, (C) molecular functions, and (D) pathway analysis of the differentially expressed genes that were suppressed by fedratinib during dibutyryl-cAMP-stimulated thermogenesis, enriched in DN-derived adipocytes [Data was cited from www.bioinformatics.com.cn/SRplot].

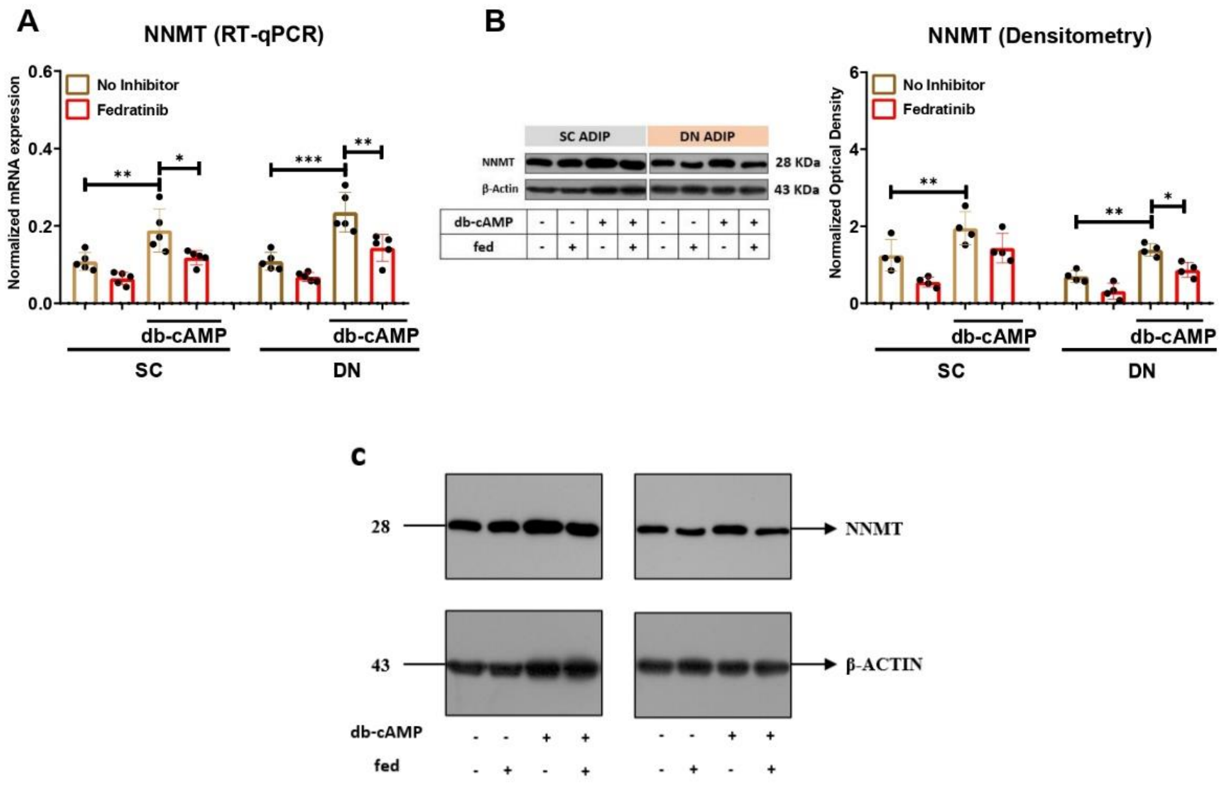


**Supplementary Figure 3. Effect of thiamine inhibition on the NNMT in subcutaneous (SC) and deep neck (DN)-derived adipocytes after 10 h of dibutyryl (db)-cAMP induced thermogenic activation.** (A) The mRNA expression of *NNMT* analyzed by RT-qPCR. (B) Protein expression of NNMT detected by immunoblotting. n=5 for RT-qPCR and n=4 for immunoblotting. Statistical analysis was performed by one-way ANOVA followed by Tukey's *post hoc* test, *p<0.05, **p<0.01, and ***p<0.001.

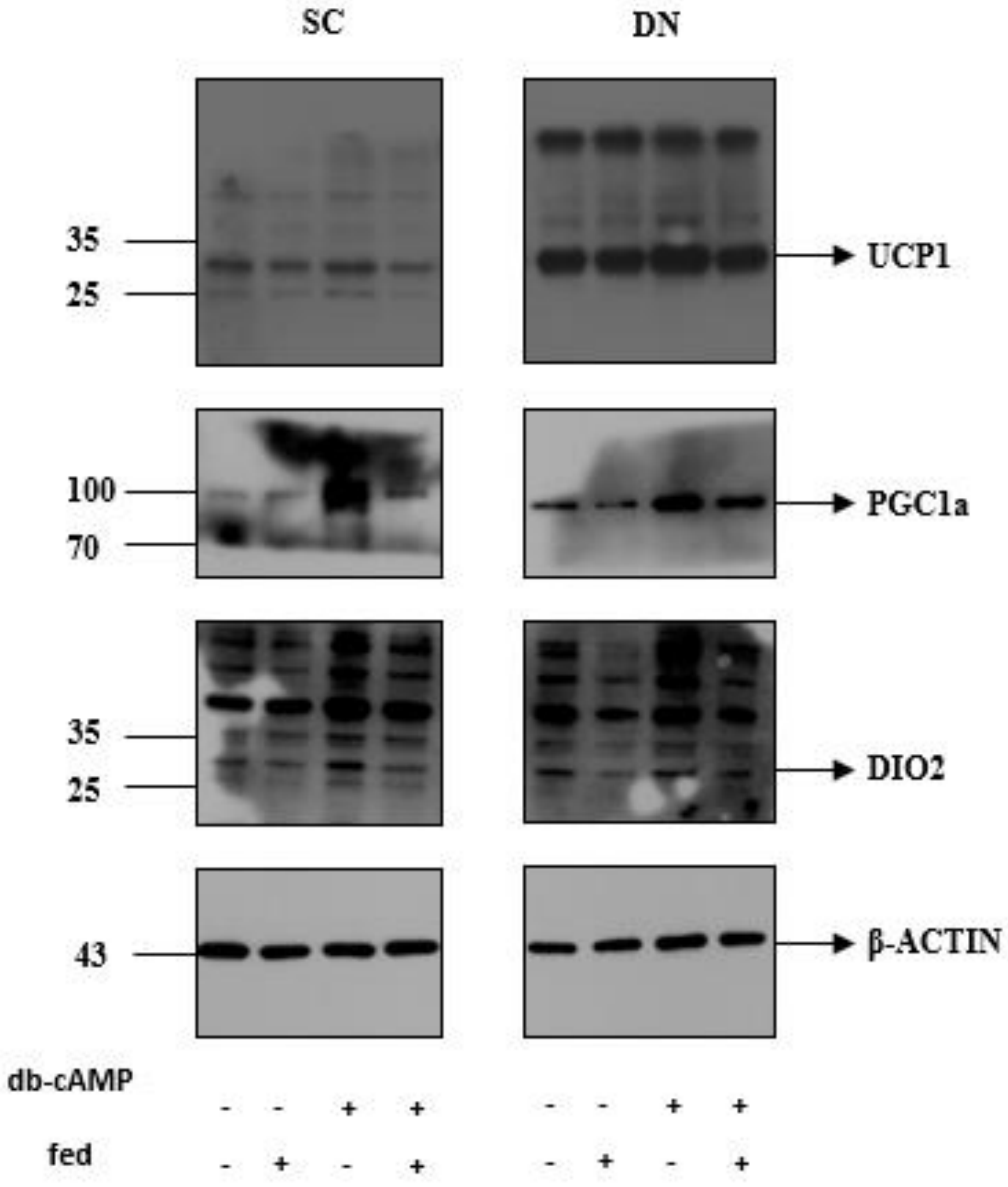


**Supplementary Figure 4. Uncropped images presented with molecular weight ladders for Figure 2B.** β-actin was used as endogenous control. Detailed information regarding the antibodies and working dilution are displayed in Supplementary Table 2.

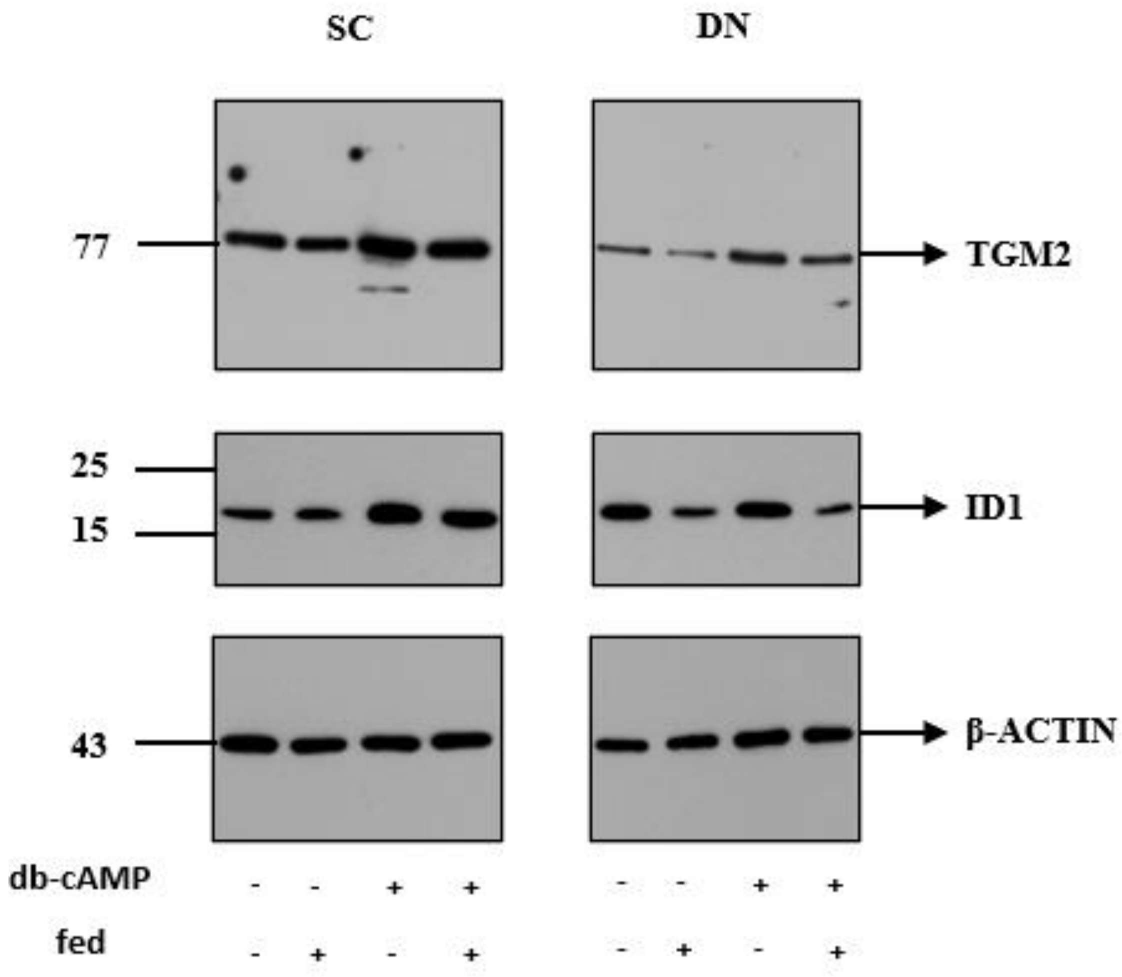


**Supplementary Figure 5. Uncropped images presented with molecular weight ladders for Figure 4B.** β-actin was used as endogenous control. Detailed information regarding the antibodies and working dilution are displayed in Supplementary Table 2.

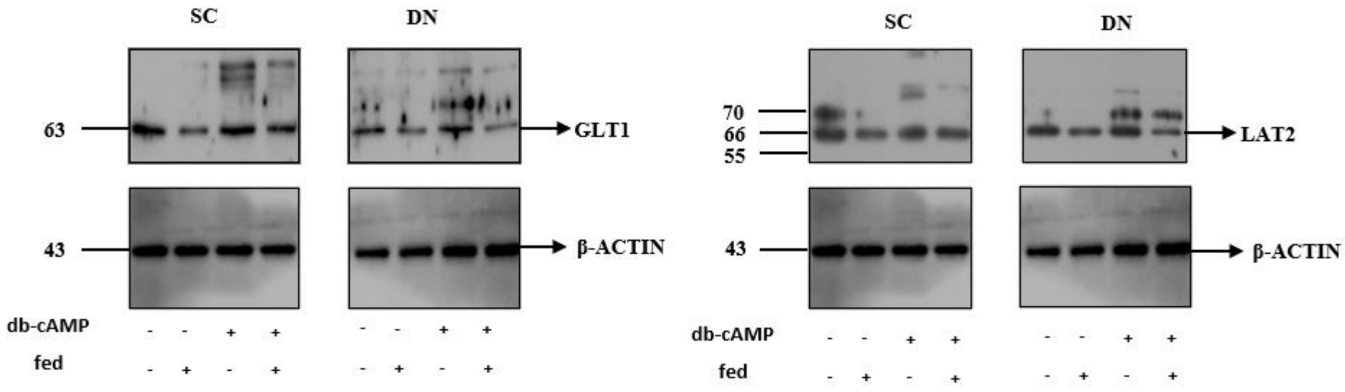


**Supplementary Figure 6. Uncropped images presented with molecular weight ladders for Figure 5D (left panel) and 5G (right panel).** β-actin was used as endogenous control. Detailed information regarding the antibodies and working dilution are displayed in Supplementary Table 2**.**

### 1.2 Supplementary Tables

**Supplementary Table 1.** Gene primers and probes

| GENES | ASSAY ID |
|---|---|
| *ACTB* | Hs01060665_g1 |
| *DIO2* | Hs00399438_m1 |
| *GAPDH* | Hs99999905_m1 |
| *ID1* | Hs00179727_m1 |
| *NNMT* | Hs00196287_m1 |
| *PGC1a* | Hs01075227_m1 |
| *SLC1A2* | Hs001102423_m1 |
| *SLC7A8* | Hs00794796_m1 |
| *TGM2* | Hs001096681_m1 |
| *UCP1* | Hs00222453_m1 |

**Supplementary Table 2.** Antibodies used in immunoblotting

| Antibody | Company | Catalog Number | Dilution |
|---|---|---|---|
| UCP1 | R&D Systems, Minneapolis, MN, USA | MAB6158 | 1:750 |
| PGC1a | Santa Cruz Biotechnology, Inc., Dallas, TX, USA | SC-517380 | 1:1000 |
| DIO2 | Thermo Fisher Scientific, Waltham, MA, USA | PA5-49631 | 1:2000 |
| TGM2 | ABCAM, Cambridge, UK | AB109200 | 1:2000 |
| NNMT | Cell Signaling Technology, Danvers, MA, USA | 24912S | 1:1000 |
| ID1 | Novus Biologicals, CO, USA | NBP2-66897 | 1:1000 |

| | | | |
|---|---|---|---|
| GLT1 | Thermo Fisher Scientific, Waltham, MA, USA | PA5-117492 | 1:1000 |
| LAT2 | Thermo Fisher Scientific, Rockville, MD, USA | TA500515 | 1:2000 |
| ACTIN | Sigma Aldrich | A2066 | 1:10000 |
| HRP-conjugated goat anti-rabbit IgG | Advansta, San Jose, CA, USA | R-05072-500 | 1:5000 |
| HRP-conjugated goat anti-mouse IgG | Advansta, San Jose, CA, USA | R-05071-500 | 1:5000 |

**Supplementary Table 3.** Differentially expressed genes (DEGs) from comparison of db-cAMP vs db-cAMP and fedratinib in subcutaneous adipocytes; n=3

| Symbol | Gene Name | Log2 Fold Change | Adjusted P value |
|---|---|---|---|
| *TGM2* | transglutaminase 2 | 2.15 | 2.33E-20 |
| *PLA2G2A* | phospholipase A2 group IIA | 1.62 | 9.73E-09 |
| *KCNK3* | potassium two pore domain channel subfamily K member 3 | 1.16 | 0.000546816 |
| *CHRDL2* | chordin like 2 | 1.15 | 0.001034331 |
| *STEAP4* | STEAP4 metalloreductase | 1.15 | 0.001150183 |
| *NNMT* | nicotinamide N-methyltransferase | 1.14 | 9.60E-05 |
| *LBP* | lipopolysaccharide binding protein | 1.13 | 0.00084268 |
| *NTRK1* | neurotrophic receptor tyrosine kinase 1 | 1.08 | 0.002846505 |
| *MGAT3* | beta-1,4-mannosyl-glycoprotein 4-beta-N-acetylglucosaminyltransferase | 1.07 | 0.002234289 |
| *ADAMTS15* | ADAM metallopeptidase with thrombospondin type 1 motif 15 | 1.06 | 0.001034331 |
| *PHYHIP* | phytanoyl-CoA 2-hydroxylase interacting protein | 1.01 | 0.008220822 |
| *TYMP* | thymidine phosphorylase | 1.00 | 0.002141746 |
| *EPSTI1* | epithelial stromal interaction 1 | 1.00 | 0.011890579 |
| *STAT4* | signal transducer and activator of transcription 4 | 0.99 | 0.010003999 |
| *SPOCD1* | SPOC domain containing 1 | 0.98 | 0.01556629 |
| *ID1* | inhibitor of DNA binding 1, HLH protein | 0.97 | 0.015646124 |
| *CCL2* | C-C motif chemokine ligand 2 | 0.95 | 0.019714157 |
| *SEMA6B* | semaphorin 6B | 0.94 | 0.023618465 |
| *LDLR* | low density lipoprotein receptor | 0.93 | 0.000850014 |
| *RIPOR3* | RIPOR family member 3 | 0.92 | 0.028817648 |
| *HSD3B7* | hydroxy-delta-5-steroid dehydrogenase, 3 beta- and steroid delta-isomerase 7 | 0.92 | 0.000850014 |
| *IL15* | interleukin 15 | 0.92 | 0.015026787 |
| *SCG2* | secretogranin II | 0.91 | 0.0345513 |
| *FER1L6* | fer-1 like family member 6 | 0.91 | 0.0345513 |
| *APOL3* | apolipoprotein L3 | 0.90 | 0.027995467 |
| *KRT7* | keratin 7 | 0.89 | 0.041862404 |
| *CCDC177* | coiled-coil domain containing 177 | 0.88 | 0.0345513 |
| *C6orf132* | chromosome 6 open reading frame 132 | 0.88 | 0.010502331 |
| *PARP9* | poly(ADP-ribose) polymerase family member 9 | 0.87 | 0.01556629 |

| | | | |
|---|---|---|---|
| *DIO2* | iodothyronine deiodinase 2 | 0.86 | 0.010318771 |
| *EGFLAM* | EGF like, fibronectin type III and laminin G domains | 0.85 | 0.0345513 |
| *SPRY1* | sprouty RTK signaling antagonist 1 | 0.84 | 0.008975406 |
| *CDC42EP4* | CDC42 effector protein 4 | 0.84 | 0.001034331 |
| *LOC730101* | uncharacterized LOC730101 | 0.83 | 0.041862404 |
| *NREP* | neuronal regeneration related protein | 0.82 | 0.007592763 |
| *RIPOR2* | RHO family interacting cell polarization regulator 2 | 0.81 | 0.020487659 |
| *LURAP1L* | leucine rich adaptor protein 1 like | 0.78 | 0.01556629 |
| *PTGS1* | prostaglandin-endoperoxide synthase 1 | 0.76 | 0.04404196 |
| *ABCC9* | ATP binding cassette subfamily C member 9 | 0.76 | 0.035580322 |
| *PTGER3* | prostaglandin E receptor 3 | 0.72 | 0.023618465 |
| *ATF4* | activating transcription factor 4 | -0.72 | 0.010533499 |
| *MT2A* | metallothionein 2A | -0.72 | 0.047898625 |
| *KIF3A* | kinesin family member 3A | -0.75 | 0.015646124 |
| *MT1X* | metallothionein 1X | -0.77 | 0.026258133 |
| *ETV5* | ETS variant transcription factor 5 | -0.77 | 0.010318771 |
| *MTHFD2* | methylenetetrahydrofolate dehydrogenase (NADP+ dependent) 2, methenyltetrahydrofolate cyclohydrolase | -0.83 | 0.015646124 |
| *SLC38A1* | solute carrier family 38 member 1 | -0.88 | 0.000466027 |
| *FOSB* | FosB proto-oncogene, AP-1 transcription factor subunit | -0.88 | 0.01556629 |
| *BIRC3* | baculoviral IAP repeat containing 3 | -0.91 | 0.0345513 |
| *HS3ST2* | heparan sulfate-glucosamine 3-sulfotransferase 2 | -0.91 | 0.0345513 |
| *ADM2* | adrenomedullin 2 | -0.92 | 0.020487659 |
| *CYP2S1* | cytochrome P450 family 2 subfamily S member 1 | -0.96 | 0.008220822 |
| *GPAT3* | glycerol-3-phosphate acyltransferase 3 | -0.97 | 0.01556629 |
| *APOLD1* | apolipoprotein L domain containing 1 | -1.17 | 3.61E-06 |
| *RRAD* | RRAD, Ras related glycolysis inhibitor and calcium channel regulator | -1.19 | 0.00076711 |

**Supplementary Table 4.** Differentially expressed genes (DEGs) from comparison of db-cAMP vs db-cAMP and fedratinib in deep neck adipocytes; n=3

| Symbol | Gene Name | Log2 Fold Change | Adjusted P value |
|---|---|---|---|
| *RIPOR2* | RHO family interacting cell polarization regulator 2 | 1.38 | 1.20E-06 |
| *SPRY1* | sprouty RTK signaling antagonist 1 | 1.37 | 2.93E-06 |
| *IL1RL1* | interleukin 1 receptor like 1 | 1.34 | 0.000173883 |
| *KCNQ3* | potassium voltage-gated channel subfamily Q member 3 | 1.28 | 0.000179194 |
| *STON1* | stonin 1 | 1.28 | 0.000739228 |
| *LYVE1* | lymphatic vessel endothelial hyaluronan receptor 1 | 1.24 | 0.001772437 |
| *ATP8B4* | ATPase phospholipid transporting 8B4 (putative) | 1.23 | 0.001032291 |
| *ARHGAP28* | Rho GTPase activating protein 28 | 1.22 | 0.00017712 |
| *LURAP1L* | leucine rich adaptor protein 1 like | 1.21 | 2.93E-06 |
| *MYPN* | myopalladin | 1.21 | 0.001109771 |
| *NNMT* | nicotinamide N-methyltransferase | 1.17 | 0.000179194 |
| *DIRAS3* | DIRAS family GTPase 3 | 1.09 | 0.015930663 |
| *CCIN* | calicin | 1.08 | 0.015362152 |
| *LXN* | latexin | 1.06 | 0.001032291 |
| *APOL3* | apolipoprotein L3 | 1.04 | 0.018921816 |
| *HSD3B7* | hydroxy-delta-5-steroid dehydrogenase, 3 beta- and steroid delta-isomerase 7 | 1.03 | 3.87E-06 |
| *BMPR1B* | bone morphogenetic protein receptor type 1B | 1.01 | 0.003292177 |
| *IGFN1* | immunoglobulin like and fibronectin type III domain containing 1 | 1.01 | 0.039038322 |
| *HGF* | hepatocyte growth factor | 1.01 | 0.031949474 |
| *CDC42EP4* | CDC42 effector protein 4 | 1.00 | 2.93E-06 |
| *PIK3IP1* | phosphoinositide-3-kinase interacting protein 1 | 1.00 | 0.038695784 |
| *LINC01605* | long intergenic non-protein coding RNA 1605 | 1.00 | 0.038173261 |
| *LMO7* | LIM domain 7 | 0.97 | 0.001065127 |
| *ZNF436* | zinc finger protein 436 | 0.95 | 0.010478386 |
| *SLC1A2* | solute carrier family 1 member 2 | 0.95 | 0.006485879 |
| *HRH1* | histamine receptor H1 | 0.95 | 0.045849513 |
| *TRAF3IP2* | TRAF3 interacting protein 2 | 0.94 | 0.003540079 |
| *SERPINB2* | serpin family B member 2 | 0.94 | 0.004616777 |
| *PHYHIP* | phytanoyl-CoA 2-hydroxylase interacting protein | 0.92 | 0.032505943 |
| *TYMP* | thymidine phosphorylase | 0.89 | 0.021333805 |
| *NOD1* | nucleotide binding oligomerization domain containing 1 | 0.89 | 0.015930663 |

| | | | |
|---|---|---|---|
| *HIF1A* | hypoxia inducible factor 1 subunit alpha | 0.88 | 0.001215782 |
| *ABI3BP* | ABI family member 3 binding protein | 0.87 | 0.006193246 |
| *KCNK2* | potassium two pore domain channel subfamily K member 2 | 0.86 | 4.97E-05 |
| *JADE2* | jade family PHD finger 2 | 0.86 | 0.004821908 |
| *ABCA1* | ATP binding cassette subfamily A member 1 | 0.86 | 0.015930663 |
| *CLEC2B* | C-type lectin domain family 2 member B | 0.83 | 0.0074486 |
| *ADAMTS5* | ADAM metallopeptidase with thrombospondin type 1 motif 5 | 0.83 | 0.003656475 |
| *ZNF423* | zinc finger protein 423 | 0.76 | 0.032505943 |
| *MCUB* | mitochondrial calcium uniporter dominant negative subunit beta | 0.69 | 0.016657911 |
| *HMGCS1* | 3-hydroxy-3-methylglutaryl-CoA synthase 1 | 0.67 | 0.001618602 |
| *ATF4* | activating transcription factor 4 | -0.58 | 0.032057653 |
| *NR4A1* | nuclear receptor subfamily 4 group A member 1 | -0.82 | 0.014001714 |
| *BAIAP2* | BAR/IMD domain containing adaptor protein 2 | -0.85 | 0.04573663 |
| *H2AC6* | H2A clustered histone 6 | -0.90 | 0.004791084 |
| *ASNS* | asparagine synthetase (glutamine-hydrolyzing) | -0.95 | 0.0199421 |
| *H4C8* | H4 clustered histone 8 | -0.99 | 0.043864342 |
| *ARC* | activity regulated cytoskeleton associated protein | -1.05 | 0.018687134 |
